# Complexity and conservation of regulatory landscapes underlie evolutionary resilience of mammalian gene expression

**DOI:** 10.1101/125435

**Authors:** Camille Berthelot, Diego Villar, Julie E. Horvath, Duncan T. Odom, Paul Flicek

## Abstract

To gain insight into how mammalian gene expression is controlled by rapidly evolving regulatory elements, we jointly analysed promoter and enhancer activity with downstream transcription levels in liver samples from twenty species. Genes associated with complex regulatory landscapes generally exhibit high expression levels that remain evolutionarily stable. While the number of regulatory elements is the key driver of transcriptional output and resilience, regulatory conservation matters: elements active across mammals most effectively stabilise gene expression. In contrast, recently-evolved enhancers typically contribute weakly, consistent with their high evolutionary plasticity. These effects are observed across the entire mammalian clade and robust to potential confounders, such as gene expression level. Overall, our results illuminate how the evolutionary stability of gene expression is profoundly entwined with both the number and conservation of surrounding promoters and enhancers.

**Highlights:**

1. Gene expression levels and stability are linked to the number of elements in the regulatory landscape.
2. Conserved regulatory elements associate with tightly controlled, highly expressed genes.
3. Recently evolved enhancers weakly influence gene expression, but promoters are similarly active regardless of conservation.
4. The interplay between complexity of the regulatory landscape and conservation of individual promoters and enhancers shapes gene expression in mammals.

## INTRODUCTION

Mammalian gene expression is controlled by collections of non-coding promoter and enhancer regions^1-3^. Numerous studies have documented the rapid evolution of mammalian regulatory elements, especially enhancers^4-9^, and yet gene expression patterns are typically highly stable between species^10-12^. How stable gene expression is maintained by rapidly evolving collections of enhancers and promoters is a fundamental question in evolutionary genetics.

Previous work connecting gene expression and regulatory evolution has typically focused on how regulatory innovations can direct lineage-specific phenotypes^4,8,13^ (reviewed in ^14^). Work across primates and mouse species has shown only limited correspondence between specific changes in gene expression and evolutionary changes in DNA methylation levels^15^, transcription factor binding^16^, or histone modifications^17^. Additionally, regulatory activities fall on a spectrum of conservation from fully conserved to lineage-specific. It is currently unknown how much insight depth of conservation provides into regulatory function partly because of a lack of datasets across divergent species^18,19^.

Here, we evaluate the consequences of regulatory evolution on gene expression divergence by jointly analyzing promoters, enhancers, and transcription levels measured in the same liver samples from over twenty mammalian species. Our results illuminate how the evolutionary resilience of gene expression is profoundly entwined with both the number and conservation of surrounding promoters and enhancers.

## RESULTS

### High conservation of gene expression levels across 25 mammals

We generated RNA sequencing (RNA-seq) data to quantify gene expression levels in liver tissue from 25 mammalian species (1-5 individuals each; Figure 1A; Methods; Supplementary Table 1). Promoters and enhancers active in liver have been reported for 20 of these species from largely the same samples^7^. Using gene annotations and orthology relationships from Ensembl^20^, we quantified and compared the gene expression for the 17,475 genes that are 1-to-1 orthologs between some or all of our study species (Figure S1; Methods).

Our results closely agree, qualitatively and quantitatively, with previous reports that tissue-specific gene expression levels are highly correlated between different mammalian species^10-12,21,22^. For ten species, analysis of the RNA-seq data was negatively affected by the relatively low quality of their reference genome assemblies (Figure 1, greyed italics; Figure S2). Although data from these species were excluded from additional analyses, we have publicly released them to allow re-analysis by the community as reference genome assemblies improve.

**Figure 1:**
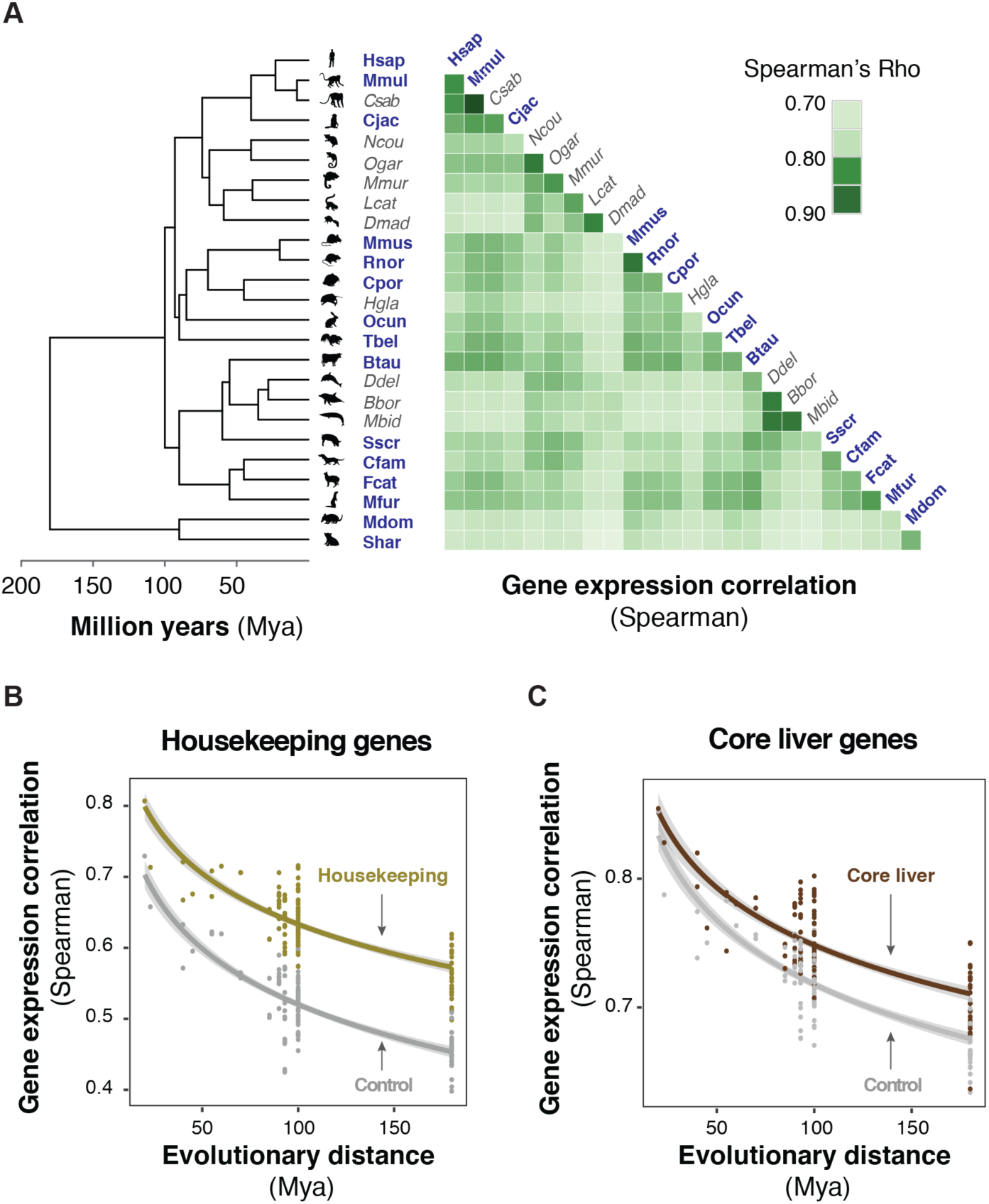
Liver gene expression levels are highly conserved across 25 mammalian species. **(A)** Pairwise correlations of normalized expression levels for 17,475 one-to-one orthologous genes in livers isolated from 25 mammalian species show high conservation of gene expression. Shading of individual tiles in the heatmap depict pairwise correlation coefficients between species (Spearman’s Rho). Known phylogenetic relationships and species divergences are represented by an evolutionary tree (left of Y-axis), which includes twenty-three placental species (in four orders) and two marsupial species (in two orders). In bolded blue: species with higher-quality reference genomes; in grey: species with either draft or proxy reference genomes (Methods). **(B-C)** Housekeeping ^23^ and core liver genes ^24^ show slower expression divergence, compared to controls. Pairwise correlation values were plotted against evolutionary distance for housekeeping (gold, **B**; 3,612 genes) and core liver genes (brown, **C**; 2,224 genes), and compared to the correlation values of control genes with the same distribution of mean expression levels across species (grey). Lines correspond to linear regression trends (after log transform of the time axis), with 95% confidence intervals in grey shading. See also Supplementary Text 1, Figure S1 and Figure S2.

We asked whether conservation of gene expression levels is higher for groups of functionally-related genes, such as broadly-expressed housekeeping genes^23^ or genes with tissue-specific liver functions^24^. Because comparing the evolutionary stability of different subsets of genes is confounded by gene expression level (Supplemental Text 1 and Figure S2), we matched each gene of interest with a control gene of similar expression, using the mean expression across species as the reference expression value (Methods). Confirming previous reports, both housekeeping and core liver gene sets exhibited higher expression correlation across species than control genes (Wilcoxon signed-rank test: both p < 2.10^-16^; Figure 1B-C)^10,25^. In addition to gene expression correlation, we also used the coefficient of variation of each gene as a measure of divergence to classify genes as evolutionarily stable or variable (i.e. inter-species standard deviation normalised by mean expression across species; Methods, Figure S2). Both housekeeping genes and core liver genes were more likely to be classified as stable (Chi-squared test: p < 2.10^-16^ and p = 2.10^-8^, respectively; Figure S2). Our results indicate that the expression levels of genes relevant to tissue function are under increased stabilizing evolutionary pressure, as proposed previously in other tissues^10,11^ and developmental contexts^26^. Nevertheless, a substantial fraction of each set was classified as variable, suggesting that functionally relevant genes can exhibit large dynamic ranges of expression across species. Thus, the coefficient of variation captures a different aspect of gene expression evolution than the expression correlation used in previous studies.

### The number of promoters and enhancers correlates with gene expression stability across evolution

We sought to connect how gene expression evolution may be directed by the evolution of promoters and enhancers in liver tissue across mammals. Specifically, we asked how gene expression levels and their evolutionary stability are affected by (i) the overall complexity of their regulatory landscape (this section), and (ii) the conservation of regulatory activity across species (next two sections). To characterize the regulatory landscape, we used the profiles of two histone modifications (H3K4me3 and H3K27ac) previously obtained using ChIP-seq in twenty mammalian species, largely from the same liver samples we report here^7^ (Table S1; Figure S3). Active promoters typically display high levels of both modifications^27,28^, whereas H3K27ac marking on its own is representative of active enhancers^29,30^ (Figure S3).

We defined the complexity of the regulatory landscape as the number of promoters and enhancers assigned to each gene in each species. As in our previous work^7^, a regulatory association domain is defined for each gene as a genomic window up and downstream of the gene’s TSS, following the strategy used by GREAT^31^ (Figure 2A, Methods). In general, this approach associates a single regulatory element to no more than two genes. Nevertheless, some gene misassignments will occur for a fraction of regulatory elements, especially among enhancers^32-34^. We use these target gene assignments to analyse the evolution of the regulatory elements associated to each of our 17,475 orthologous genes.

**Figure 2:**
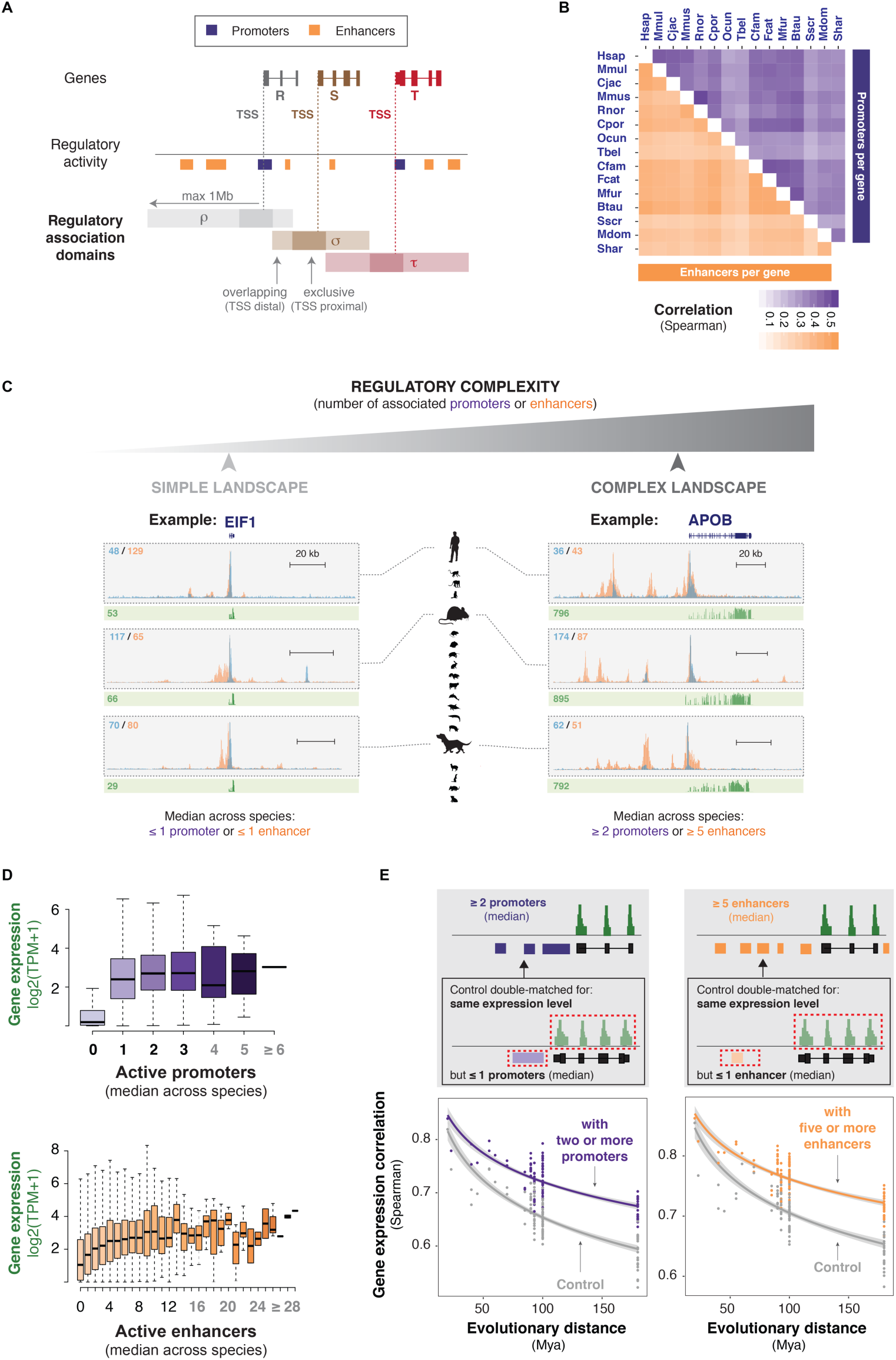
The number of promoters and enhancers corresponds with gene expression stability across evolution. **(A)** Genes are associated with all active regulatory elements sitting between their TSS and the TSS of the next gene on either side, within a limit of 1Mb. Regulatory elements sitting directly on the TSS of a gene (max. 5kb upstream and 1kb downstream) were exclusively associated to that gene (darker shading, exclusive TSS proximal). The cartoon example illustrates this procedure for three genes R, S, and T; the active regulatory elements in their vicinity (regulatory activity) and the resulting regulatory association domains ρ, σ and τ. **(B)** The number of promoters and enhancers associated to a gene is correlated across species. Shading of individual tiles in the heatmap corresponds to pairwise tie-corrected Spearman correlation coefficients for numbers of promoters and enhancers associated to orthologous genes across 15 mammalian species. **(C)** Examples of genes with simple (*EIF1*) and complex (*APOB*) regulatory landscapes in liver. Regulatory complexity was measured as the median number of promoters and enhancers associated to each gene across species. Genomic tracks are displayed for three representative species (human, mouse and dog): histone modification ChIP-seq fold enrichments are shown in blue (H3K4me3) and orange (H3K27ac), and RNA-seq reads in green. Numbers in blue and orange correspond to the maximum fold enrichment for each histone mark; numbers in green correspond to the gene expression values in TPM normalised across species. **(D)** Gene expression distributions (mean expression across species) are shown for genes associated with increasing numbers of active promoters (purple) or active enhancers (orange) in an average mammal. The number of active enhancers associated to a gene has an additive effect, whereas promoter activity shows a more switch-like effect on gene expression levels. Classes containing fewer than thirty genes are greyed on the x axis. **(E)** The number of promoters and enhancers associated per gene contributes to evolutionary stability of gene expression. **Grey insets**: Gene expression divergence across species is compared between (i) genes associated to multiple promoters or enhancers (top) and (ii) control genes with the same expression level but associated to few promoters or enhancers (one or none, bottom). **Plots**: Pairwise Spearman correlation coefficients of expression levels between species were plotted against evolutionary distance for genes associated with multiple promoters (left; 1,688 genes) or enhancers (right; 1,479 genes), and compared to control gene sets. In both cases the number of associated promoters or enhancers corresponds to the median number across species. Lines are as described in Figure 1B-C. See also Figure S3, Figure S4 and Figure S5.

We observed that the regulatory complexity is moderately correlated across species (Figure 2B), reflecting the known rapid evolution of mammalian regulatory elements^7^. To summarise the regulatory landscape at each gene, we took the median number of promoters and enhancers across species as a representative value in an average mammal (Figure 2C). Genes associated with larger numbers of transcriptional regulatory elements tend to be more highly expressed (Figure 2D and S4), agreeing with observations in a single species^35-37^. This was especially true for enhancers, suggesting that the majority of the active enhancers identified have a measurable effect on gene expression (Figure 2D). Conversely, promoters appear to have more of a switch-like effect, where at least one active promoter is necessary to turn the gene on, but additional promoters are not associated with substantially higher gene expression levels. The associations we observe were not due to biased ChIP-seq signal intensity or artefacts associated with highly expressed genes (Figure S4), or the method of defining regulatory association domains (Figure S5).

We next asked whether the number of promoters or enhancers associated to a gene also influence the evolutionary conservation of gene expression levels. To do this, genes associated to multiple promoters or enhancers across species were compared to control genes matched for expression level but with only a simple regulatory landscape (Figure 2E, grey insets). Genes with regulatory inputs from multiple promoter or enhancer elements showed significantly increased expression conservation (Wilcoxon signed-rank test: promoters and enhancers both p < 2.10^-16^; Figure 2E).

Overall, these observations support a direct connection among the complexity of the regulatory landscape, gene expression, and gene expression conservation. In the remaining sections, we leverage our extensive phylogenetic scope to explore how regulatory conservation and regulatory complexity together influence gene expression evolution.

### Conserved regulatory activity associates with high and evolutionarily stable gene expression

Active promoters are largely functionally conserved across mammalian species, while enhancer activity evolves rapidly^6,7,9,13,38^. Conserved regulatory regions are thought to be particularly important for gene expression control^39-41^, but definitive evidence for this beyond individual cases is generally lacking^18,42,43^.

In previous work, we identified a set of 1872 promoters and 279 enhancers that exhibit conserved activity in the livers of most placental mammals (“placental-conserved” regulatory elements, Figure 3A; Figure S3)^7^. Placental-conserved elements typically are a minority within a gene’s regulatory landscape, although these conserved regulatory elements may disproportionally contribute to the levels and/or stability of gene expression.

**Figure 3:**
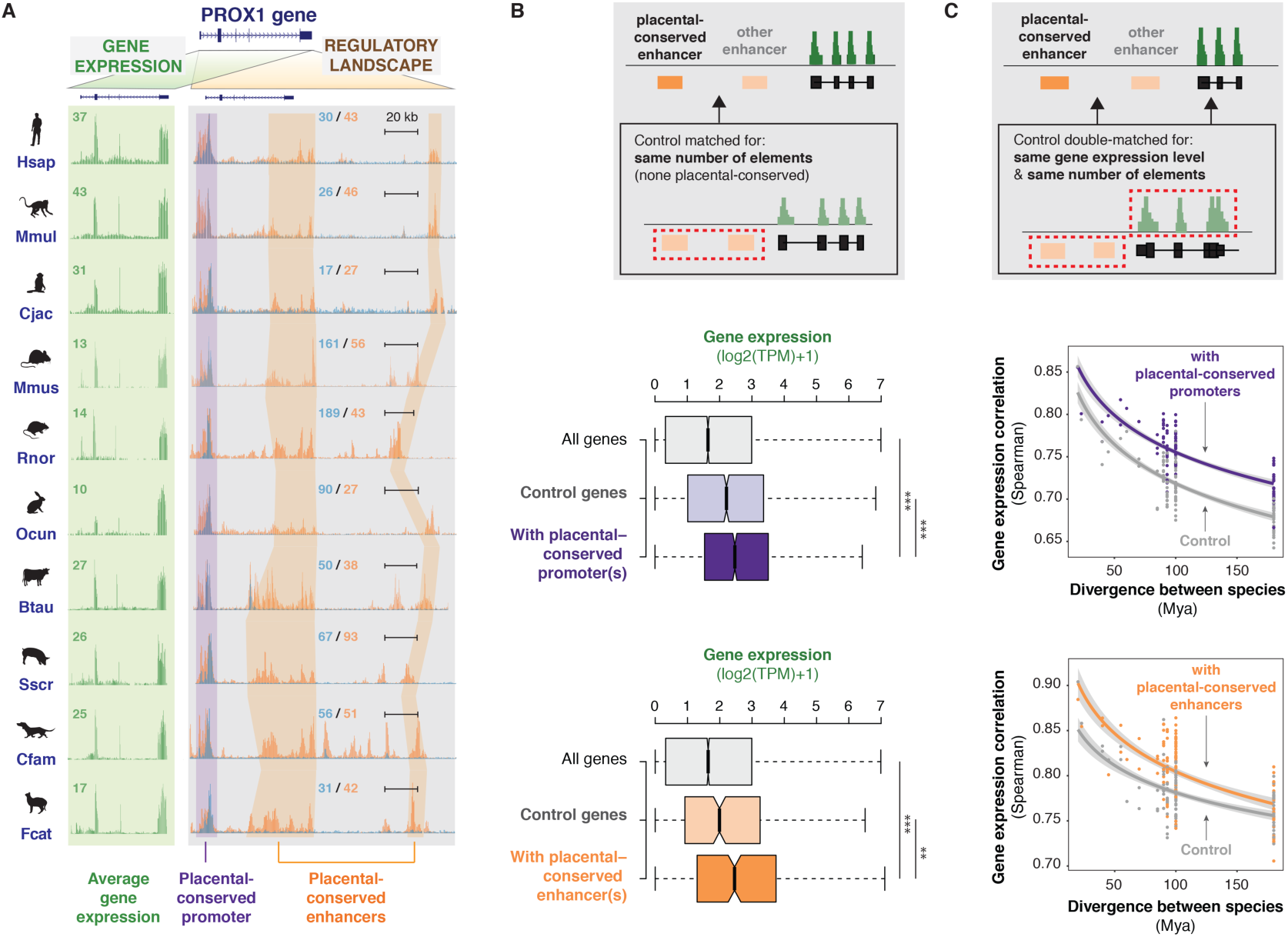
Conserved regulatory activity is associated with both high and stable gene expression levels. **(A)** Example of gene expression and regulatory landscapes around the PROX1 gene in livers from ten placental mammals. Each row shows PROX1 expression (left, green background) and activity of promoters and enhancers around the PROX1 locus in one species (H3K4me3 (blue) and H3K27ac (orange) ChIP-seq signals, grey background; as described in Figure 2C). A placental-conserved promoter and two placental-conserved enhancers at this locus are highlighted. **(B)** Genes associated with placental-conserved promoters and enhancers show high expression levels. **Grey inset:** The contribution of placental-conserved regulatory activity to gene expression was evaluated using control genes associated with the same number of active promoters or enhancers, none of which are placental-conserved. **Boxplots** show the distribution of mean expression levels across species for all 1-to-1 orthologs (all genes); for genes associated with placental-conserved elements (dark purple for promoters, 2,384 genes; dark orange for enhancers, 387 genes); and for control genes (pale purple for promoters, pale orange for enhancers). ***: p < 0.001, **: p < 0.01, Wilcoxon rank sum test. **(C)** Genes associated to placental-conserved promoters and enhancers exhibit slow expression divergence across species. **Grey inset:** The contribution of placental-conserved regulatory activity to gene expression conservation was evaluated using control genes with similar expression levels and associated with the same number of active promoters or enhancers, none of which are placental-conserved. **Plots**: Pairwise Spearman correlation coefficients of expression levels between species were plotted against evolutionary distance, for genes associated with placental-conserved promoter(s) (purple) or enhancer(s) (orange) and control gene sets. Lines are as described in Figure 1B-C. See also Figure S3 and Figure S6.

We first asked whether placental-conserved regulatory elements contribute more to gene expression levels than other elements. Genes associated with conserved elements exhibited higher transcription levels than control genes associated with the same number of regulatory elements where none are placental-conserved (Figure 3B; Wilcoxon rank sum test: promoters p = 2.10^-8^; enhancers p = 0.001). This result was consistent whether the expression was measured using the mean expression across all species or in a representative species (e.g. human; Figure S6). Thus, highly expressed genes appear to be associated with regulatory regions more likely to be maintained during evolution. Indeed, housekeeping and core liver genes are significantly more likely to be associated with placental-conserved promoters (Figure S6).

We next isolated the contribution of placental-conserved regulatory activity to gene expression stability. We compared sets of genes matched for expression levels and total number of associated regulatory regions, but differing by the presence or absence of placental-conserved elements. Genes associated with placental-conserved elements were more correlated in expression across species than those without (Figure 3C; Wilcoxon signed-rank test: promoters p < 2.10^-16^; enhancers p = 7.10^-16^). Analyzing expression stability based on the coefficient of variation further supports the enhanced importance of conserved regulatory elements (Figure S6).

Taken together, these results demonstrate that deeply conserved elements contribute disproportionately to maintaining both high and stable gene expression levels across species.

### Recently-evolved regulatory activity has modest impact on gene expression levels

We also previously identified a set of 794 promoters and 10,434 enhancers that were reproducibly active in human liver, but not active in the liver of any other of our study species (Figure 4A; Figure S3 and Methods). By combining these with our newly obtained gene expression data, we were able to ask whether recently-evolved regulatory elements influence gene expression (Figure 4B), and if so how much (Figure 4C).

**Figure 4:**
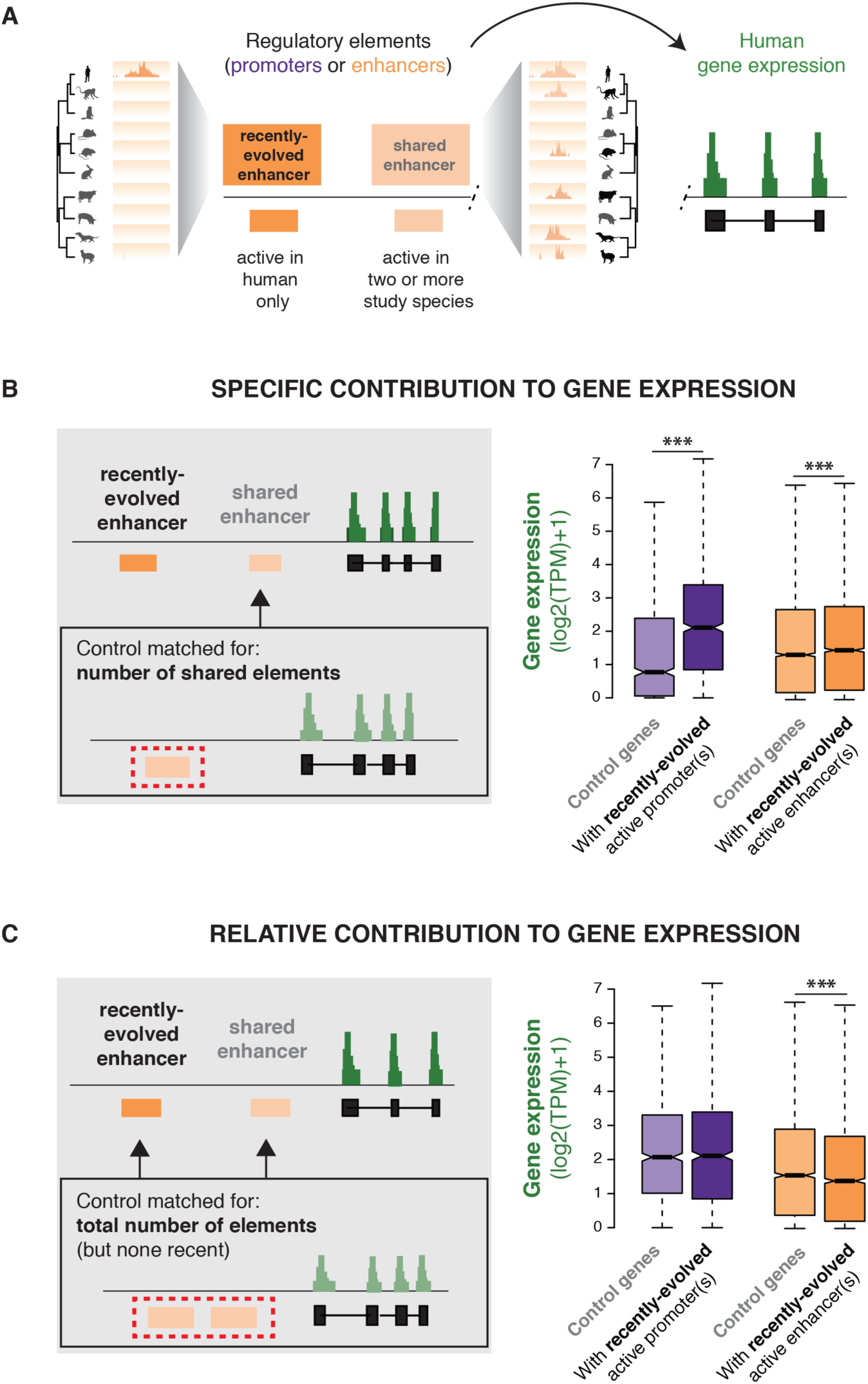
Recently-evolved enhancer activities weakly contribute to gene expression levels. **(A)** The contribution of recently-evolved regulatory elements (active in a single study species, here human) to gene expression was analysed. Genes with recently-evolved regulatory elements are typically also associated with shared regulatory elements (active in two or more species). **(B)** When compared to control genes with the same number of shared regulatory elements, human genes associated with additional recently-evolved promoter(s) or enhancer(s) exhibit significantly higher expression levels (***: p < 0.001, Wilcoxon rank sum test; promoters: 995 matched genes; enhancers: 5,173 matched genes). **(C)** When compared to control genes with the same total number of regulatory elements, human genes associated with recently-evolved enhancer(s) (orange) are expressed at lower levels (***: p < 0.001, Wilcoxon rank sum test; 3,054 matched genes). Recently evolved promoters are as active as shared ones (purple; 995 matched genes). See also Figure S3, and Figure S7 for similar results in other species.

The human genes putatively regulated by recently-evolved promoters are typically expressed well above background and show no difference in expression compared to control genes with more conserved promoters (Wilcoxon rank sum test: p = 0.64; Figure 4C). New promoters therefore seem as likely to be functional as those shared with at least one other species: indeed, 57% of genes targeted by a recently-evolved promoter in human apparently did not rely on any other promoter for expression in liver.

Whether recently-evolved enhancers have a measurable effect on gene expression is appreciably more problematic to establish, largely because identifying enough control genes was challenging. Specifically, 42% of human genes with 1-to-1 orthologs across placentals are associated with recently-evolved enhancers, and these genes were more likely to be associated with enhancers shared across several species (mean: 3.3 vs. 0.8 shared enhancers; Wilcoxon rank sum test: p < 2.10^-16^). We therefore limited our analyses to the subsets of human genes that could be matched for expression level and/or landscape complexity (Methods).

Overall, our results revealed that recently-evolved enhancers typically increase gene expression slightly less than do shared enhancers. First, the presence of recently-evolved enhancer(s) is associated with modestly higher expression, when compared to control genes with the same background of evolutionarily shared enhancers, (Wilcoxon rank sum test: p = 4.10^-5^; Figure 4B). Second, compared to genes with the same total number of enhancers, genes with recently-evolved enhancer(s) exhibit slightly lower expression (Wilcoxon rank sum test: p = 2.10^-6^; Figure 4C). These results were also observed in other species (Figure S7).

Together, these observations suggest that recently-evolved regulatory elements have a measurable effect on gene expression. Our results paint a picture of recently-evolved enhancers as weaker than those active in several different species, yet they are at least partly functional and pervasively modulate gene expression across species.

### Recently-evolved elements consistently contribute to increased expression stability

We asked if recently-evolved regulatory activity also has a weaker stabilising impact on gene expression than do shared regulatory elements. At the scale of a single species (human), we observed that genes with and without recently-evolved regulatory elements showed no difference in expression conservation when controlling for total number of regulatory elements and expression level (Wilcoxon signed-rank test: promoters p = 0.43, enhancers p = 0.24; Figure S8). Moreover, recently-evolved human enhancers were equally likely to be associated to genes with either evolutionarily stable or variable expression (Chi-squared test: p = 0.11); if anything, recently-evolved promoters weakly associated with stable genes (Chi-squared test: p = 0.03). Thus, recently-evolved regulatory activity in a single species has no obvious relationship with expression divergence between species.

Interestingly, recently-evolved promoters and enhancers from different species concentrated more often than expected in the vicinity of the same genes (Figure 5A-C). This effect remained significant regardless of the size of the regulatory association domain (Figure S8). We delineated a set of genes that recurrently associate with recently-evolved elements across different mammals (Figure 5A; Methods). Surprisingly, these genes were significantly more correlated in expression than expected based on their expression levels (Wilcoxon signed-rank test: promoters and enhancers both p < 2.10^-16^; Figure 5D). However, these genes also exhibited particularly complex regulatory landscapes (1.3 active promoters and 8.6 active enhancers on average), which associate with stable gene expression. When controlling for both expression level and the total number of enhancers, genes with a recurrent accumulation of recently-evolved enhancers exhibited faster divergence in their expression levels than those without (Wilcoxon signed-rank test: promoters and enhancers both p < 2.10^-16^; Figure 5E). In contrast, the accumulation of recently-evolved promoters associated with increased gene expression stability.

**Figure 5:**
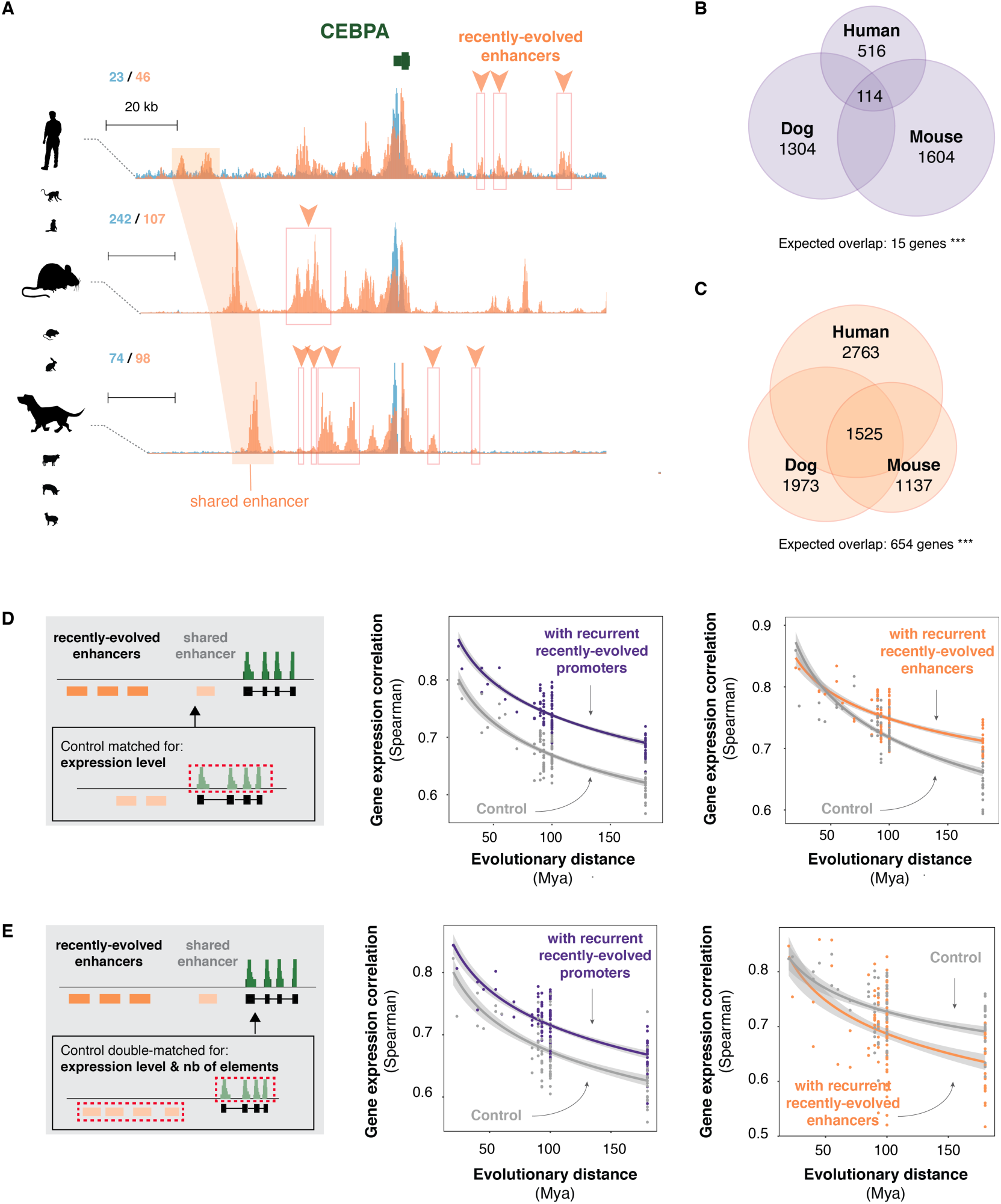
Multiple recently-evolved regulatory elements contribute to gene expression stability. **(A)** Example of recurrent association of a gene with recently-evolved enhancers in multiple species. Genomic tracks show the regulatory landscape around the liver-specific gene *CEBPA* in human, mouse and dog (H3K4me3 (blue) and H3K27ac (orange) ChIP-seq signals; as described in Figure 2C). Recently-evolved enhancer activity in the three species is delineated with orange boxes and arrowheads. An orthologous enhancer with conserved activity across species is highlighted with orange shading. **(B-C)** Genes associated with recently-evolved regulatory activity significantly overlap across three reference species (**B**: promoters; **C**: enhancers; ***: p < 0.001, Chi-squared test). Numbers in Venn diagrams correspond to the number of genes with recently-evolved elements in all three species (center) and restricted to a single species. Overlaps between pairs of species are not shown. **(D)** Genes recurrently associated with recently-evolved elements across species exhibit high conservation of expression. Pairwise Spearman correlation coefficients of expression levels between species were plotted against evolutionary distance for genes recurrently associated with recently-evolved promoters (purple; 1,208 matched genes) or enhancers (orange; 729 matched genes) across multiple species, and control genes with similar mean expression levels across species. Lines are as described in Figure 1B-C. **(E)** Compared to control genes with similar expression levels and regulatory complexity, genes associated with recurrent recently-acquired promoter activity in multiple species diverge more slowly in expression (purple; 1,207 matched genes). Recently-evolved enhancers however are weaker at stabilising gene expression evolution: genes recurrently associated with recently-evolved enhancers across species exhibit higher divergence than control genes with similar expression levels and number of enhancers (orange; 207 matched genes). Plots as above. See also Figure S8.

These results suggest that recently-evolved elements contribute to gene expression stability across species by maintaining the complexity of the regulatory landscape. Nevertheless recently-evolved enhancers appear weaker at buffering expression changes than do elements conserved across several species.

### The composite liver regulatory landscape across mammals

Previous sections have rigorously quantified the impact of the regulatory elements that are either conserved across species (Figure 3), or singular to one species in the dataset (Figures 4 and 5); however, these elements make up less than half of the regulatory regions identified in every species (Figure S3). Here, we exploit the full extent of our genome-wide datasets to characterize the continuous relationship between regulatory evolution and gene expression.

We built a reference-free map of the regulatory landscape across mammalian species (Methods; Figure 6A) by projecting all twenty regulatory landscapes onto a single summary landscape for each orthologous gene, to create 17,475 meta-genes. These meta-genes collect all the independent regulatory elements associated to a gene, regardless of the number or subset of species in which each element is active (Figure 6A). Therefore each metagene’s summary landscape explicitly integrates both regulatory complexity and regulatory conservation. The reference-free map treats each meta-promoter and meta-enhancer as a single evolutionary acquisition and describes regulatory evolution with simple metrics (total number of accumulated elements across lineages and number of species where the activity is present; Methods). On average, meta-genes were associated with 2.3 active meta-promoters and 11 active meta-enhancers (sd: 2.2 and 13.0, respectively).

**Figure 6:**
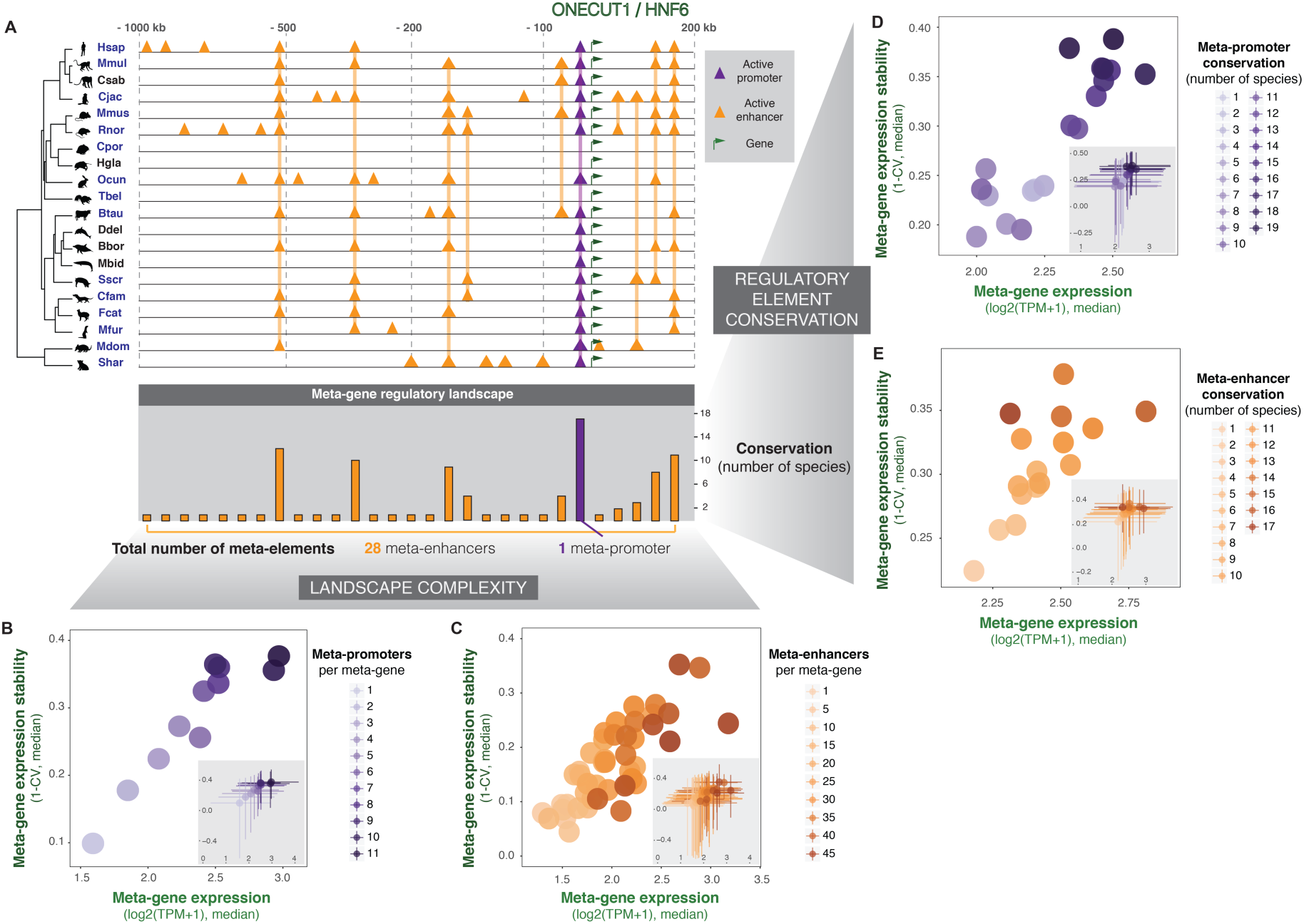
An integrated summary of the evolution of mammalian regulatory complexity. **(A)** Representative example of the reference-free approach to connect promoter and enhancer activity with gene expression across species. Tracks in each of twenty species show an indicative landscape of active promoters and enhancers around the ONECUT1 gene, with orthologous regions linked across species by vertical lines. The reference-free mapping of these regulatory elements across species results in a meta-gene regulatory landscape that includes a single meta-promoter and 28 meta-enhancers (bottom barplot, x-axis). For each meta-element, evolutionary conservation is recorded as the number of species where promoter or enhancer activity is detected (y-axis). **(B-C)** The number of meta-promoters (**B**, purple) or meta-enhancers (**C**, orange) in the meta-gene landscape correlates with increased expression levels (x-axis) and expression stability (y-axis). Meta-genes were categorised according to the number of meta-promoters or meta-enhancers in their regulatory landscape. For each category, the median gene expression level is plotted against the median expression stability (1-CV, where CV—coefficient of variation across species). Insets in each plot show the spread of the distributions (interquartile ranges). Classes containing fewer than 30 meta-genes are not shown. **(D-E)** The evolutionary conservation of meta-promoters (**D**, purple) and meta-enhancers (**E**, orange) correlates with increasingly high and stable gene expression. Individual meta-promoters or meta-enhancers were classified according to the number of species where their activity is detected, and the median expression levels and expression stability of their putative target meta-genes were plotted as above. Insets as above. Classes containing fewer than 30 meta-genes are not plotted.

To investigate the overall impact of regulatory complexity on gene expression evolution, we stratified meta-genes by their number of associated meta-promoters (Figure 6B) and meta-enhancers (Figure 6C). We observed that expression level and expression stability increase steadily with regulatory complexity across the entire mammalian gene set. Interestingly, this trend was consistent for promoters as well as enhancers, whereas in an average mammal, promoters show a more switch-like behaviour (Figure 2). Therefore, integrating regulatory information across twenty species increases the resolution to detect the impact of multiple meta-promoters.

We next asked how the entire spectrum of regulatory conservation impacts gene expression across the full set 17,475 orthologous genes. Meta-promoters (Figure 6D) and meta-enhancers (Figure 6E) were classified by the number of species in which they are active, and we measured the expression and evolutionary stability of their associated meta-genes. Strikingly, the level and stability of gene expression continuously tracks with the conservation of the regulatory landscape. This result extends and complements our observations on highly-conserved and recently-evolved elements, and suggests that the gradual relationship between conservation of regulatory landscapes and stability of expression is a general feature of mammalian gene regulation.

These data also illustrate the difficulty of predicting expression level or expression stability of specific genes, even when informed by enhancer and promoter maps from twenty mammals. In fact, we observe substantial variability within, and overlap between, all of the meta-element classes (insets of Figure 6B-E). Nevertheless, our integrated analysis reveals how regulatory complexity and conservation interplay to shape expression level and expression stability across mammalian genomes.

## DISCUSSION

To date, comparative approaches to understanding gene regulation have largely focused on lineage-specific regulatory innovations, thus identifying candidate regions driving lineage-specific phenotypes^4,8,13,15^ (reviewed in ^14^). At the same time, evolutionarily conserved regulatory elements are thought to play a predominant role in gene regulation^40,44,45^, while the functional impact and broader role of regulatory elements with low signals of conservation has been the subject of speculation^46-48^. To extend these analyses, we collected and analysed an integrated dataset of gene expression output and regulatory histone marks from the same liver samples across 20 mammalian species. This strategy allowed us to systematically test the contributions of both landscape complexity (i.e. number of regulatory elements) and the conservation of promoters and enhancers on gene expression evolution. Our methodology simultaneously evaluates regulatory activities that range from completely-conserved across mammals to present in only one species.

Our key finding is that the transcription of a gene is evolutionarily stabilised by the presence of many regulatory elements regardless of the conservation of those elements. In other words, gene expression level and its evolutionary resilience reflect the complexity of the regulatory landscape, both within a single species and across mammals. However, regulatory regions are not functionally equal: those highly-conserved across placental mammals exert a more powerful stabilizing effect, associating with gene expression levels that are simultaneously high and evolutionarily stable. In contrast, recently-evolved enhancers contribute more weakly to gene expression and transcriptional stability, consistent with a model whereby a fraction of newborn enhancer elements have a neutral role on gene expression evolution^49,50^. These effects are clear throughout our data, whether considering a full-scale, reference-free map of mammalian regulatory complexity, or investigating subsets of extremely conserved or divergent regulatory elements. Our discoveries extend previous reports connecting evolutionary constraint on promoter and enhancer activities with expression outputs^4,16,17,51^, and are consistent with an enhanced functional importance of conserved regulatory elements^39,41,52^.

There are a number of limitations to our approach. First, the precise measurement of regulatory complexity, conservation, and gene expression are partly dependent on the reference genome assembly and annotation, both of which are known to be of variable completeness across our study species. Second, our strategy to connect regulatory elements to putative functional targets is based on genomic proximity, which may mis-assign distal enhancers; this could explain in part the fuzzy correlation between enhancer activity and gene expression. Experimentally linking regulatory activity to target genes would enable a more accurate evaluation of how individual enhancer elements contribute to transcriptional output^53-55^. Third, mapping regulatory activity in other tissues or additional signatures of regulatory activity, such as open chromatin^56^, other histone modifications^57^ or co-activator proteins^58^, may identify other features that contribute to gene expression evolution. Fourth, we did not explore how the often-poorly annotated non-coding transcriptome evolves, where different regulatory principles may apply^59,60^. Regardless, as experimental technologies and genome references improve, the datasets we report here will become increasingly valuable.

This study suggests a general framework of how transcriptional output and transcriptional regulation co-evolve. Active regulatory elements have long been known to additively contribute to gene expression control^39,61^. By investigating transcriptional control in twenty species and connecting this to gene expression, we demonstrate how the number of active promoters and enhancers relates to gene expression stability across mammals. Our evolutionary analysis supports recent reports indicating that synergistic interactions are rare among enhancer clusters^2,62,63^. Our observations are consistent with existing models of enhancer function^64-67^, and provide mechanistic insight into how conserved transcriptional outputs can be achieved by complex and rapidly evolving regulatory landscapes.

## METHODS

### Ethics statement

The investigation was approved by the Animal Welfare and Ethics Review Board and followed the Cambridge Institute guidelines for the use of animals in experimental studies under Home Office license PPL 70/7535. Human liver samples were obtained under Human Tissue Act license 08-H0308-117 from the Addenbrooke’s Hospital at the University of Cambridge with patients’ consent, and was approved by the National Research Ethics Service.

### Source and detail of tissues

We quantified gene expression profiles in liver samples from 25 species by RNA extraction coupled to high throughput sequencing (RNA-seq), typically from 2-4 individuals per species. In most cases, these are the same samples we previously used in ChIP-seq experiments to assess regulatory activity across twenty mammals^7^. The origin, number of replicates, sex and age for each species’ samples are detailed in Table S1.

### Total RNA-sequencing (RNA-seq) library preparation

Total RNA was extracted from snap-frozen liver tissue with RNAeasy Mini Kit (Qiagen). 20 mg of tissue were weighed on dry-ice and immediately homogenized in 600 microliters (ul) of RLT buffer containing 10 ul of beta-mercaptoethanol per mililiter of buffer. Tissue samples were homogenized in a Precellys 24 tissue homogenizer, using settings 5500-2x15-015 and Precellys tubes CK28-R (bertin technology). Liver homogenates were processed according to manufacturers’ instructions (Qiagen RNAeasy Mini Kit) and total RNA eluted in 50ul RNAse-free water. 10 ug total RNA from each sample were treated with 4 units of Turbo DNAse (Ambion), and total RNA samples were run on an Agilent Bioanalyser (RNA nano chip) to check RNA integrity. Samples were taken forward if RIN values were above 7. Ribosomal RNA (rRNA) was depleted with Ribo-Zero rRNA removal kit (Epicentre RZC110424) as per instructions from the manufacturer, using 5 ug of DNAse-treated total RNA.

Strand-specific rRNA-depleted RNA-seq libraries were prepared with a modified version of TruSeq RNA Library Preparation kit (Illumina). Fragmentation and first-strand synthesis of rRNA-depleted RNA samples were according to the Illumina protocol. Second-strand cDNA synthesis was done with SuperScript double-stranded cDNA synthesis kit (Life Technologies) at 16C for two hours, using a 10mM dATP, dCTP, dGTP, dUTP nucleotide mix. cDNA was purified with QIAquick PCR purification kit (Qiagen) and end repair, A-tailing and adaptor ligation were performed with Illumina’s protocol. Second-strand degradation was achieved by treatment with one unit of Uracil N-Glycosylase (Life Technologies) at 35C for 15 minutes, prior to PCR enrichment. Libraries were amplified according to Illumina’s protocol for 13 PCR cycles, and cleaned-up with Agencourt AMPure beads (Beckman Coulter) with a 1:1 DNA:beads ratio. RNA-seq libraries were quantified with Kapa Library quantification kit (Kapa biosystems) on a QuantStudio 6 Flex instrument (Applied Biosystems), pooled in equimolar amounts and sequenced to a minimum depth of twenty million uniquely mapped reads on an Illumina HiSeq 2500 instrument. Libraries were sequenced as either paired-end 100 bp or paired-end 150bp.

### RNA-seq alignment and gene expression quantification

RNA-seq reads from de-multiplexed fastq files were trimmed to 100 bp and aligned to the corresponding reference genomes and full transcript sets available in Ensembl and Ensembl Pre! v.73^20^ using TopHat 2.0.13^68^ with default parameters and a mate pair inner distance (-r) of 75 bp. Aligned reads were subsampled to 20 million read pairs per sample.

Transcript quantification was performed using Cufflinks 2.2.1^69^ (default parameters), based on the transcript annotations available in Ensembl v.73^20^. Estimated gene expression levels in FPKM (fragment per kb of exon per million mapped fragments) were obtained from the Cufflinks gene summary output. The gene expression levels were further transformed into TPM (transcripts per million transcripts).

Genes annotated in human with 1-to-1 orthologs in one or more species were identified from the gene phylogenies available in Ensembl v.73^20^. In each species where a unique orthologue was identified, the mean expression level over all available replicates was used as the representative expression level for this gene. Orthologous expression levels were further normalized between species using the median of ratios to the geometric means, as described in ^70^.

### Assignment of active regulatory regions to putative target genes

Regulatory elements were assigned to putative target genes following rules similar to those implemented in GREAT^31^. A regulatory association domain was defined for each gene as the genomic window up and downstream of the TSS, until the TSS of the next gene and within 1 Mb. Additionally, genes were exclusively assigned those regulatory elements directly at the TSS (up to 5 kb upstream and 1 kb downsteam). In general, this approach associates a single regulatory element to no more than two genes, with a few exceptions in case of overlapping genes and/or extremely close TSSs. For each gene, the TSS annotation used was that of the reference (“canonical”) transcript in Ensembl v.73^20^.

This procedure was performed in each species independently. The median number of promoters or enhancers assigned to each orthologous gene across species was used as the representative value for this gene in an average mammal.

### Measures of gene expression divergence

Evolutionary divergence of gene expression was measured by the Spearman correlation coefficients for orthologous expression levels between pairs of species. The relative divergence of two gene subsets was compared using the correlations within each subset across all pairs of species (Wilcoxon paired rank sum test). Species phylogenetic trees were built by hierarchical clustering based on the pairwise correlations of gene expression levels using complete linkage. The unweighted pair group method with arithmetic mean (UPGMA) gave similar results. When estimating the relative divergence of particular gene subsets, confounding effects due to differences in expression levels distributions were controlled for by matching genes one-to-one to control genes with similar expression. Control matching was performed in R with the MatchIt library^71^ using the caliper option to prune genes that could not be matched to an appropriate control (increments of 0.1 to 0.001). Matching for the number of regulatory elements was performed similarly, either on its own or in combination with gene expression level.

Expressed genes (mean expression across species > 1 TPM; 10,704 genes) were also classified into evolutionarily stable and variable genes based on their coefficient of variation (standard deviation across species normalized by mean expression; bottom 50%: stable; top 50%: variable). Because these two categories had different mean expression levels (18.8 vs. 27.5 TPM; Wilcoxon rank sum test: p < 2.10^-16^), we additionally matched stable and variable genes into pairs with similar mean expression across species as described above with MatchIt (4,264 genes in each group), and removed 2,176 unmatched genes from the subsets.

### Recently-evolved regulatory elements

Recently-evolved regulatory elements were identified in each of the ten highest-quality reference genomes in our dataset (human, macaque, marmoset, mouse, rat, rabbit, cow, pig, dog and cat), all of which are included in the multiple whole genome alignment available from Ensembl. Regulatory elements were defined as recently-evolved when they either could not be aligned to an orthologous sequence in any of the other genomes, or when their orthologous loci in other genomes showed no significant enrichment in regulatory histone marks, as described in ^7^. As previously reported, the vast majority of recently-evolved promoters corresponded to non-alignable sequences. Most recently-evolved enhancers could be aligned to orthologous loci in other species, but these orthologous locations showed no evidence of regulatory activity^7^.

Genes recurrently targeted by recently-evolved promoters were defined as genes associated with a recently-evolved promoter in five species or more out of the ten (i.e. median number of recently-evolved promoters across species ≥ 0.5; 1,075 genes). Genes recurrently targeted by recently-evolved enhancers were defined as genes associated with three or more recently-evolved enhancers in more than five species (i.e. median number of recently-evolved enhancers across species ≥ 3; 530 genes). Conversely, control genes rarely targeted by recently-evolved promoters were defined as genes associated with no recently-evolved promoter in five species or more; genes rarely targeted by recently-evolved enhancers had one or no recently-evolved enhancers in five species or more.

### All-vs-all inter-species analysis of promoter and enhancer activity

Regulatory elements identified in each species were first mapped to their orthologous loci in each of the ten highest-quality reference genomes in our dataset (human, macaque, marmoset, mouse, rat, rabbit, cow, pig, dog and cat). These ten species are all cross-mappable against each other via a single multi-species alignment. Elements from the other species are mapped via pairwise alignments to one or more of these ten reference species, as described in ^7^. In each of these ten coordinate systems, regulatory elements active in two or more species were considered to be orthologous and merged into a consensus element when their genomic coordinates overlapped by 50% or more, using the *bedmap* utility from BEDOPS v.2.4.20 (option ‐‐fraction-either 0.5)^72^. This procedure resulted in ten independent maps of regulatory activity; each integrating regulatory elements from the 10 or more species aligned to this reference. All ten independent maps were then merged into a master regulatory map containing meta-elements. This map thereby integrates all regulatory elements identified in all species as long as a given element had an orthologous locus in at least one of the ten reference species.

Meta-elements in the master regulatory map were assigned to putative target meta-genes based on the collection of predicted targets in each individual species. Specifically, a gene was considered a predicted target if it was identified as such in at least half of the species where the regulatory element is active. Similarly, meta-elements were identified as meta-promoters or meta-enhancers based on their predominant histone marking across the orthologous elements integrated in the map.

## Data availability

RNA-sequencing data has been deposited under Array Express accession number E-MTAB-4550, with the exception of three human and four mouse datasets (previously reported in E-MTAB-4052). ChIP-seq data from twenty mammalian species were previously reported in ArrayExpress accession number E-MTAB-2633. Python and R scripts used to process the data are available upon request.

## AUTHOR CONTRIBUTIONS

CB, DV, PF, DTO designed experiments; DV performed experiments; CB, DV analyzed the data; JEH provided tissue samples; CB, DV, PF, DTO wrote the manuscript; PF, DTO oversaw the work. All authors read and approved the final manuscript.

## ACKNOWLEDGEMENTS

We thank the Cambridge Institute Biological Resource Unit and Genomics Core for technical support, Claudia Kutter, Christine Feig, Maša Roller and Tim Rayner for useful comments and discussions, and the EMBL-EBI systems team for management of computational resources. This research was supported by Cancer Research UK grant 20412; the European Research Council grant 615584; the Welcome Trust grant numbers WT108749/Z/15/Z, WT098051, WT202878/A/16/Z and WT202878/B/16/Z; the French Institut National de la Santé et de la Recherche Médicale (INSERM) and the European Molecular Biology Laboratory. Cetacean samples were collected by the UK Cetacean Strandings Investigation Programme, funded by Defra and the Governments of Scotland and Wales. The authors declare no conflicts of interest.

## REFERENCES

1 Spitz, F. & Furlong, E. E. Transcription factors: from enhancer binding to developmental control. Nature reviews. Genetics 13, 613–626, doi:10.1038/nrg3207 (2012).

2 Moorthy, S. D. et al. Enhancers and super-enhancers have an equivalent regulatory role in embryonic stem cells through regulation of single or multiple genes. Genome research 27, 246–258, doi:10.1101/gr.210930.116 (2017).

3 Shin, H. Y. et al. Hierarchy within the mammary STAT5-driven Wap super-enhancer. Nature genetics 48, 904–911, doi:10.1038/ng.3606 (2016).

4 Cotney, J. et al. The evolution of lineage-specific regulatory activities in the human embryonic limb. Cell 154, 185–196, doi:10.1016/j.cell.2013.05.056 (2013).

5 Xiao, S. et al. Comparative epigenomic annotation of regulatory DNA. Cell 149, 1381–1392, doi:S0092-8674(12)00574-0 [pii] 10.1016/j.cell.2012.04.029 (2012).

6 Vierstra, J. et al. Mouse regulatory DNA landscapes reveal global principles of cis-regulatory evolution. Science 346, 1007–1012, doi:10.1126/science.1246426 (2014).

7 Villar, D. et al. Enhancer evolution across 20 mammalian species. Cell 160, 554–566, doi:10.1016/j.cell.2015.01.006 (2015).

8 Reilly, S. K. et al. Evolutionary genomics. Evolutionary changes in promoter and enhancer activity during human corticogenesis. Science 347, 1155–1159, doi:10.1126/science.1260943 (2015).

9 Young, R. S. et al. The frequent evolutionary birth and death of functional promoters in mouse and human. Genome research 25, 1546–1557, doi:10.1101/gr.190546.115 (2015).

10 Brawand, D. et al. The evolution of gene expression levels in mammalian organs. Nature 478, 343–348, doi:10.1038/nature10532 (2011).

11 Chan, E. T. et al. Conservation of core gene expression in vertebrate tissues. J Biol 8, 33, doi:jbiol130 [pii] 10.1186/jbiol130 (2009).

12 Merkin, J., Russell, C., Chen, P. & Burge, C. B. Evolutionary dynamics of gene and isoform regulation in Mammalian tissues. Science 338, 1593–1599, doi:10.1126/science.1228186 (2012).

13 Prescott, S. L. et al. Enhancer divergence and cis-regulatory evolution in the human and chimp neural crest. Cell 163, 68–83, doi:10.1016/j.cell.2015.08.036 (2015).

14 Reilly, S. K. & Noonan, J. P. Evolution of Gene Regulation in Humans. Annual review of genomics and human genetics 17, 45–67, doi:10.1146/annurev-genom-090314-045935 (2016).

15 Pai, A. A., Bell, J. T., Marioni, J. C., Pritchard, J. K. & Gilad, Y. A genome-wide study of DNA methylation patterns and gene expression levels in multiple human and chimpanzee tissues. PLoS genetics 7, e1001316, doi:10.1371/journal.pgen.1001316 (2011).

16 Wong, E. S. et al. Decoupling of evolutionary changes in transcription factor binding and gene expression in mammals. Genome research 25, 167–178, doi:10.1101/gr.177840.114 (2015).

17 Cain, C. E., Blekhman, R., Marioni, J. C. & Gilad, Y. Gene expression differences among primates are associated with changes in a histone epigenetic modification. Genetics 187, 1225–1234, doi:10.1534/genetics.110.126177 (2011).

18 Chatterjee, S., Bourque, G. & Lufkin, T. Conserved and non-conserved enhancers direct tissue specific transcription in ancient germ layer specific developmental control genes. BMC developmental biology 11, 63, doi:10.1186/1471-213X-11-63 (2011).

19 Necsulea, A. & Kaessmann, H. Evolutionary dynamics of coding and non-coding transcriptomes. Nature reviews. Genetics 15, 734–748, doi:10.1038/nrg3802 (2014).

20 Flicek, P. et al. Ensembl 2013. Nucleic acids research 41, D48–55, doi:10.1093/nar/gks1236 (2013).

21 Sudmant, P. H., Alexis, M. S. & Burge, C. B. Meta-analysis of RNA-seq expression data across species, tissues and studies. Genome biology 16, 287, doi:10.1186/s13059-015-0853-4 (2015).

22 Perry, G. H. et al. Comparative RNA sequencing reveals substantial genetic variation in endangered primates. Genome research 22, 602–610, doi:10.1101/gr.130468.111 (2012).

23 Eisenberg, E. & Levanon, E. Y. Human housekeeping genes, revisited. Trends in genetics : TIG 29, 569–574, doi:10.1016/j.tig.2013.05.010 (2013).

24 Odom, D. T. et al. Control of pancreas and liver gene expression by HNF transcription factors. Science 303, 1378–1381, doi:10.1126/science.1089769 (2004).

25 She, X. et al. Definition, conservation and epigenetics of housekeeping and tissue-enriched genes. BMC genomics 10, 269, doi:10.1186/1471-2164-10-269 (2009).

26 Israel, J. W. et al. Comparative Developmental Transcriptomics Reveals Rewiring of a Highly Conserved Gene Regulatory Network during a Major Life History Switch in the Sea Urchin Genus Heliocidaris. PLoS biology 14, e1002391, doi:10.1371/journal.pbio.1002391 (2016).

27 Santos-Rosa, H. et al. Active genes are tri-methylated at K4 of histone H3. Nature 419, 407–411, doi:10.1038/nature01080 (2002).

28 Heintzman, N. D. et al. Distinct and predictive chromatin signatures of transcriptional promoters and enhancers in the human genome. Nature genetics 39, 311–318, doi:10.1038/ng1966 (2007).

29 Creyghton, M. P. et al. Histone H3K27ac separates active from poised enhancers and predicts developmental state. Proceedings of the National Academy of Sciences of the United States of America 107, 21931–21936, doi:10.1073/pnas.1016071107 (2010).

30 Rada-Iglesias, A. et al. A unique chromatin signature uncovers early developmental enhancers in humans. Nature 470, 279–283, doi:10.1038/nature09692 (2011).

31 McLean, C. Y. et al. GREAT improves functional interpretation of cis-regulatory regions. Nature biotechnology 28, 495–501, doi:10.1038/nbt.1630 (2010).

32 Whalen, S., Truty, R. M. & Pollard, K. S. Enhancer-promoter interactions are encoded by complex genomic signatures on looping chromatin. Nature genetics 48, 488–496, doi:10.1038/ng.3539 (2016).

33 Sanyal, A., Lajoie, B. R., Jain, G. & Dekker, J. The long-range interaction landscape of gene promoters. Nature 489, 109–113, doi:10.1038/nature11279 (2012).

34 Sikora-Wohlfeld, W., Ackermann, M., Christodoulou, E. G., Singaravelu, K. & Beyer, A. Assessing computational methods for transcription factor target gene identification based on ChIP-seq data. PLoS computational biology 9, e1003342, doi:10.1371/journal.pcbi.1003342 (2013).

35 Odom, D. T. et al. Core transcriptional regulatory circuitry in human hepatocytes. Molecular systems biology 2, 2006 0017, doi:10.1038/msb4100059 (2006).

36 Mikkelsen, T. S. et al. Comparative epigenomic analysis of murine and human adipogenesis. Cell 143, 156–169, doi:10.1016/j.cell.2010.09.006 (2010).

37 Arnold, C. D. et al. Genome-wide quantitative enhancer activity maps identified by STARR-seq. Science 339, 1074–1077, doi:10.1126/science.1232542 (2013).

38 Jubb, A. W., Young, R. S., Hume, D. A. & Bickmore, W. A. Enhancer Turnover Is Associated with a Divergent Transcriptional Response to Glucocorticoid in Mouse and Human Macrophages. Journal of immunology 196, 813–822, doi:10.4049/jimmunol.1502009 (2016).

39 Cheng, Y. et al. Principles of regulatory information conservation between mouse and human. Nature 515, 371–375, doi:10.1038/nature13985 (2014).

40 Lindblad-Toh, K. et al. A high-resolution map of human evolutionary constraint using 29 mammals. Nature 478, 476–482, doi:10.1038/nature10530 (2011).

41 McLean, C. & Bejerano, G. Dispensability of mammalian DNA. Genome research 18, 1743–1751, doi:10.1101/gr.080184.108 (2008).

42 Kvon, E. Z. et al. Progressive Loss of Function in a Limb Enhancer during Snake Evolution. Cell 167, 633–642 e611, doi:10.1016/j.cell.2016.09.028 (2016).

43 Royo, J. L. et al. Transphyletic conservation of developmental regulatory state in animal evolution. Proceedings of the National Academy of Sciences of the United States of America 108, 14186–14191, doi:10.1073/pnas.1109037108 (2011).

44 Siepel, A. et al. Evolutionarily conserved elements in vertebrate, insect, worm, and yeast genomes. Genome research 15, 1034–1050, doi:10.1101/gr.3715005 (2005).

45 Wittkopp, P. J. & Kalay, G. Cis-regulatory elements: molecular mechanisms and evolutionary processes underlying divergence. Nature reviews. Genetics 13, 59–69, doi:10.1038/nrg3095 (2011).

46 Arnold, C. D. et al. Quantitative genome-wide enhancer activity maps for five Drosophila species show functional enhancer conservation and turnover during cis-regulatory evolution. Nature genetics 46, 685–692, doi:10.1038/ng.3009 (2014).

47 Lowdon, R. F., Jang, H. S. & Wang, T. Evolution of Epigenetic Regulation in Vertebrate Genomes. Trends in genetics : TIG 32, 269–283, doi:10.1016/j.tig.2016.03.001 (2016).

48 Sundaram, V. et al. Widespread contribution of transposable elements to the innovation of gene regulatory networks. Genome research 24, 1963–1976, doi:10.1101/gr.168872.113 (2014).

49 Cooper, G. M. & Brown, C. D. Qualifying the relationship between sequence conservation and molecular function. Genome research 18, 201–205, doi:10.1101/gr.7205808 (2008).

50 Kellis, M. et al. Defining functional DNA elements in the human genome. Proceedings of the National Academy of Sciences of the United States of America 111, 6131–6138, doi:10.1073/pnas.1318948111 (2014).

51 Zhou, X. et al. Epigenetic modifications are associated with inter-species gene expression variation in primates. Genome biology 15, 547, doi:10.1186/s13059-014-0547-3 (2014).

52 Villar, D., Flicek, P. & Odom, D. T. Evolution of transcription factor binding in metazoans - mechanisms and functional implications. Nature reviews. Genetics 15, 221–233, doi:10.1038/nrg3481 (2014).

53 Mifsud, B. et al. Mapping long-range promoter contacts in human cells with high-resolution capture Hi-C. Nature genetics 47, 598–606, doi:10.1038/ng.3286 (2015).

54 Schoenfelder, S. et al. The pluripotent regulatory circuitry connecting promoters to their long-range interacting elements. Genome research 25, 582–597, doi:10.1101/gr.185272.114 (2015).

55 Kieffer-Kwon, K. R. et al. Interactome maps of mouse gene regulatory domains reveal basic principles of transcriptional regulation. Cell 155, 1507–1520, doi:10.1016/j.cell.2013.11.039 (2013).

56 Buenrostro, J. D., Giresi, P. G., Zaba, L. C., Chang, H. Y. & Greenleaf, W. J. Transposition of native chromatin for fast and sensitive epigenomic profiling of open chromatin, DNA-binding proteins and nucleosome position. Nature methods 10, 1213–1218, doi:10.1038/nmeth.2688 (2013).

57 Pradeepa, M. M. et al. Histone H3 globular domain acetylation identifies a new class of enhancers. Nature genetics 48, 681–686, doi:10.1038/ng.3550 (2016).

58 May, D. et al. Large-scale discovery of enhancers from human heart tissue. Nature genetics 44, 89–93, doi:10.1038/ng.1006 (2011).

59 Necsulea, A. et al. The evolution of lncRNA repertoires and expression patterns in tetrapods. Nature 505, 635–640, doi:10.1038/nature12943 (2014).

60 Kutter, C. et al. Rapid turnover of long noncoding RNAs and the evolution of gene expression. PLoS genetics 8, e1002841, doi:10.1371/journal.pgen.1002841 (2012).

61 Visel, A. et al. Functional autonomy of distant-acting human enhancers. Genomics 93, 509–513, doi:10.1016/j.ygeno.2009.02.002 (2009).

62 Bothma, J. P. et al. Enhancer additivity and non-additivity are determined by enhancer strength in the Drosophila embryo. eLife 4, doi:10.7554/eLife.07956 (2015).

63 Hay, D. et al. Genetic dissection of the alpha-globin super-enhancer in vivo. Nature genetics 48, 895–903, doi:10.1038/ng.3605 (2016).

64 Arnosti, D. N. & Kulkarni, M. M. Transcriptional enhancers: Intelligent enhanceosomes or flexible billboards? Journal of cellular biochemistry 94, 890–898, doi:10.1002/jcb.20352 (2005).

65 Biggin, M. D. Animal transcription networks as highly connected, quantitative continua. Developmental cell 21, 611–626, doi:10.1016/j.devcel.2011.09.008 (2011).

66 Long, H. K., Prescott, S. L. & Wysocka, J. Ever-Changing Landscapes: Transcriptional Enhancers in Development and Evolution. Cell 167, 1170–1187, doi:10.1016/j.cell.2016.09.018 (2016).

67 Panne, D. The enhanceosome. Current opinion in structural biology 18, 236–242, doi:10.1016/j.sbi.2007.12.002 (2008).

68 Kim, D. et al. TopHat2: accurate alignment of transcriptomes in the presence of insertions, deletions and gene fusions. Genome biology 14, R36, doi:10.1186/gb-2013-14-4-r36 (2013).

69 Trapnell, C. et al. Differential gene and transcript expression analysis of RNA-seq experiments with TopHat and Cufflinks. Nature protocols 7, 562–578, doi:10.1038/nprot.2012.016 (2012).

70 Anders, S. & Huber, W. Differential expression analysis for sequence count data. Genome biology 11, R106, doi:10.1186/gb-2010-11-10-r106 (2010).

71 Ho, D., Imai, K., King, G. & Stuart, E. A. MatchIt: Nonparametric Preprocessing for Parametric Causal Inference. Journal of Statistical Software 42, 1–28 (2011).

72 Neph, S. et al. BEDOPS: high-performance genomic feature operations. Bioinformatics 28, 1919–1920, doi:10.1093/bioinformatics/bts277 (2012).

